# Pathogenic BRCA1 DBD variants exhibit altered DNA binding affinities and susceptibility to menadione

**DOI:** 10.1101/2025.07.10.664210

**Authors:** Emma Cismas, Esme Lowry, Vereena Salib, Kaitlin Lowran, Colin G. Wu

**Affiliations:** Department of Chemistry, Oakland University, Rochester, MI, 48309, USA

## Abstract

Breast Cancer Susceptibility Gene 1 (*BRCA1*) codes for a DNA repair protein that facilitates the repair of double-stranded DNA breaks (DSBs) in human cells through the homologous recombination (HR) pathway. Mutations of *BRCA1* are highly associated with breast cancer; however, many variants remain unclassified with unknown cellular phenotypes. The DNA binding activity of BRCA1 is localized primarily to its central region, which can be divided into two distinct domains: DNA Binding Domain 1 (DBD1; amino acids (aa) 330-554) and 2 (DBD2; aa 894-1057). We previously proposed a model in which DBD1 targets BRCA1 to DSBs for the promotion of DNA end resection, while DBD2 targets BRCA1 to telomeres to function in chromatin remodeling and telomere regulation. In this study, we hypothesized that unknown DBD variants (T374I, K408E, N417S, N909I, M1008I, and R1028H) with similar properties to known disease-causing variants (Q356H, F461L, R496H, D940Y, S1027N, and E1038G) would also be pathogenic. The affinities of each variant for single-stranded DNA (ssDNA), double-stranded DNA (dsDNA), and a G-quadruplex (G4) sequence were measured via biolayer interferometry. The DNA repair phenotypes of each variant were analyzed by overexpression in HEK cells to determine correlation between binding activity and DNA damage response. Altogether, these results provide insight into how missense mutations affect the ability of BRCA1 DBDs to facilitate the DNA damage response.

## INTRODUCTION

Breast Cancer Susceptibility Gene 1 (*BRCA1*) codes for a human tumor suppressor protein that was discovered in 1990 (1). BRCA1 has since been identified to play a role in DNA double-stranded break (DSB) repair, telomere maintenance, and the repair and restart of stalled DNA replication forks (2–6). Replication fork stress is often induced by DNA damage, fragile sites, or the presence of non-canonical DNA (7); if left unmanaged, stress can result in DSBs and lead to tumorigenesis (8, 9). Numerous repair mechanisms may be employed to alleviate this stress, such as homologous recombination (HR), non-homologous end joining, alternative end joining, and single-strand annealing (10–13). Similarly, non-canonical DNA structures occur naturally and must be accurately managed throughout DNA repair and replication processes. BRCA1 preferentially binds to non-canonical forms of DNA (14–17) and plays a role in the regulation of telomeres. For example, G-quadruplex (G4) DNA is a form of non-canonical DNA that is abundant in regions with high levels of genome regulation. If left to accumulate, G4s can stall DNA repair and replication (18–20). With a general sequence of (GGGN_1-7_)_4_ (with N being any nucleotide), over 400,000 human G4 motifs have been identified (21).

Most studies have focused on the interaction of BRCA1 with other proteins through its N-terminal RING domain (aa 1-109) and C-terminal BRCT repeats (aa 1646-1863). For example, BRCA1 RING forms a heterodimer with BARD1 to act as a tumor suppressor complex that participates in HR (22). The BRCT domain of BRCA1 is instead homologous to domains found in other DNA repair proteins (23), and can bind DNA or proteins such as FANCJ to promote replication fork restart at G4 DNA (24–27). In contrast, the central region of BRCA1 (aa 452-1057) is largely disordered with few globular domains, yet flexible to allow for interaction with both DNA substrates and partner proteins such as RAD51 and PALB2 (28, 29). Due to its ability to directly bind DNA, this central region is often referred to as the DNA Binding Domain (DBD) (14). The DBD can be further studied as two separate domains that bind DNA with higher affinity than other fragments within the central region: DBD1 (aa 330-554) and DBD2 (aa 894-1057) (14, 16, 17, 28, 30–32). Little other structural information about the DBDs is known as most studies have focused on the characterization of the RING and C-terminal BRCT repeats (33, 34).

Previous work has shown that DBD1 preferentially binds dsDNA, while DBD2 has a preference for ssDNA and G4 DNA (35). This suggests that the DBDs specialize in targeting BRCA1 to either DSBs (for the promotion of DNA end resection) or to telomeres (for general regulation), respectively. According to structural predictions, the DBDs are located on opposing sides of the solvent-accessible surfaces of BRCA1 (Figure 1), which allows for this separation of tasks.

**Figure 1.**
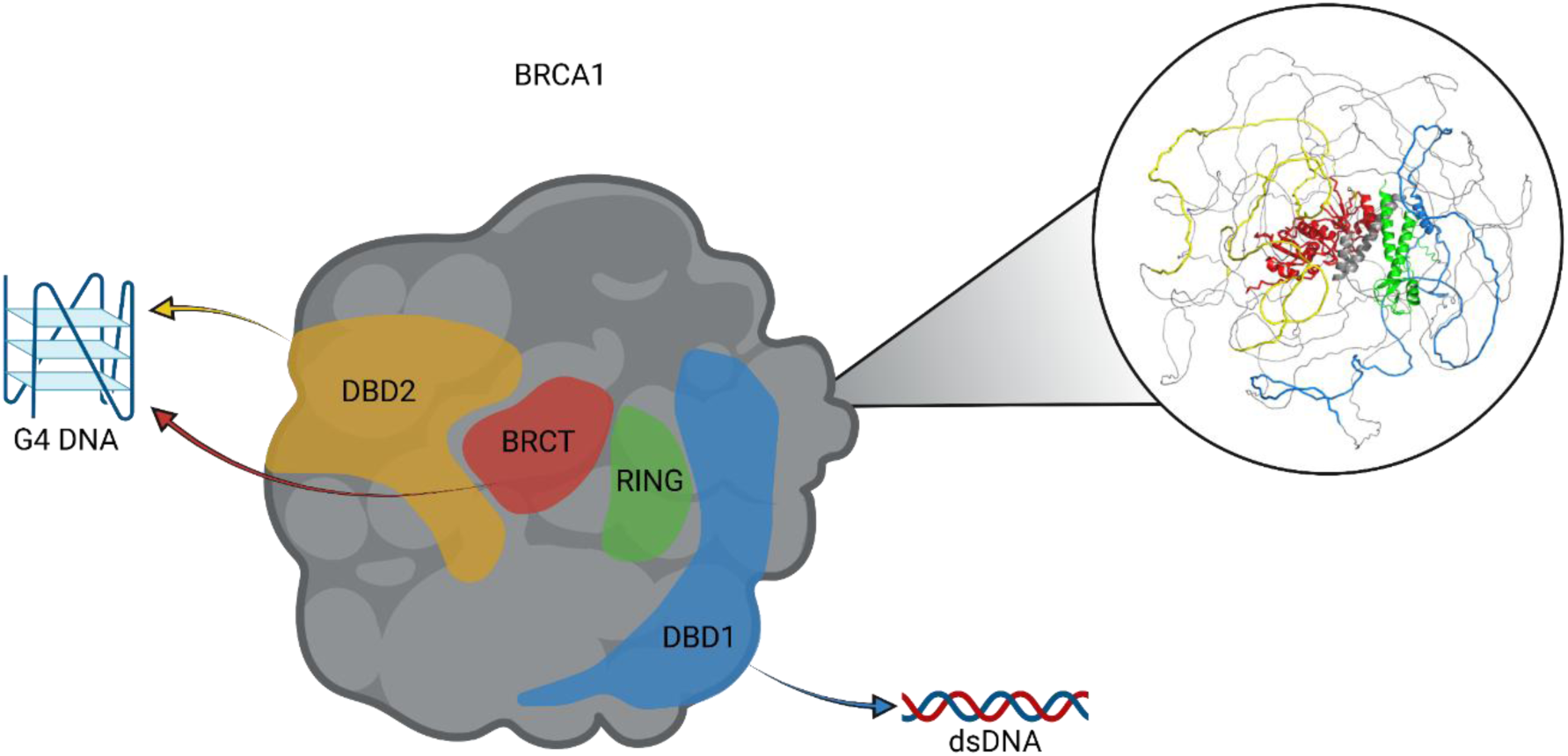
Cartoon and structural prediction of full-length BRCA1 generated in AlphaFold 2. Highlighted domains include RING (aa 1-109, green), DBD1 (aa 330-554, blue), DBD2 (aa 894-1057, yellow), and the tandem BRCT repeats (aa 1646-1863, red). Color mapping corresponds with Figure 2.

The Leiden Open Variation Database (LOVD) stores reported *BRCA1* mutations for collaboration among researchers and clinicians and has identified over 1800 mutations in human *BRCA1* (36). These mutations are implicated in 52% of familial breast and ovarian cancer cases (37), and are also associated with the development of cardiovascular disease due to its crucial role in cellular repair (38–40). Patients with *BRCA1*-related breast cancers specifically experience a significant increase in the incidence of breast cancer, an earlier onset, a higher relapse rate, and a more aggressive disease. While the LOVD has identified cancerous mutations in *BRCA1*, there remain numerous variants of unknown significance (VUS) with unknown phenotypic effects. This study examines the DNA binding affinities of twelve DBD variants (Figure 2), six of which are classified as pathogenic (DBD1 Q356H, R496H, and F461L; DBD2 D940Y, S1027N, and E1038G), and six of which are unclassified VUSs (DBD1 T374I, K408E, and N417S; DBD2 N909I, M1008I, and R1028H). Binding interactions were determined for single-stranded (ss), double-stranded (ds), and G4 DNA as these types of DNA appear as intermediates in the DNA repair processes discussed previously. These results are compared to cellular assays of BRCA1 overexpressed in HEK cells, current LOVD predictions, and structural predictions generated in AlphaFold 2 to provide insight into the pathogenicity of these unclassified VUSs.

**Figure 2.**
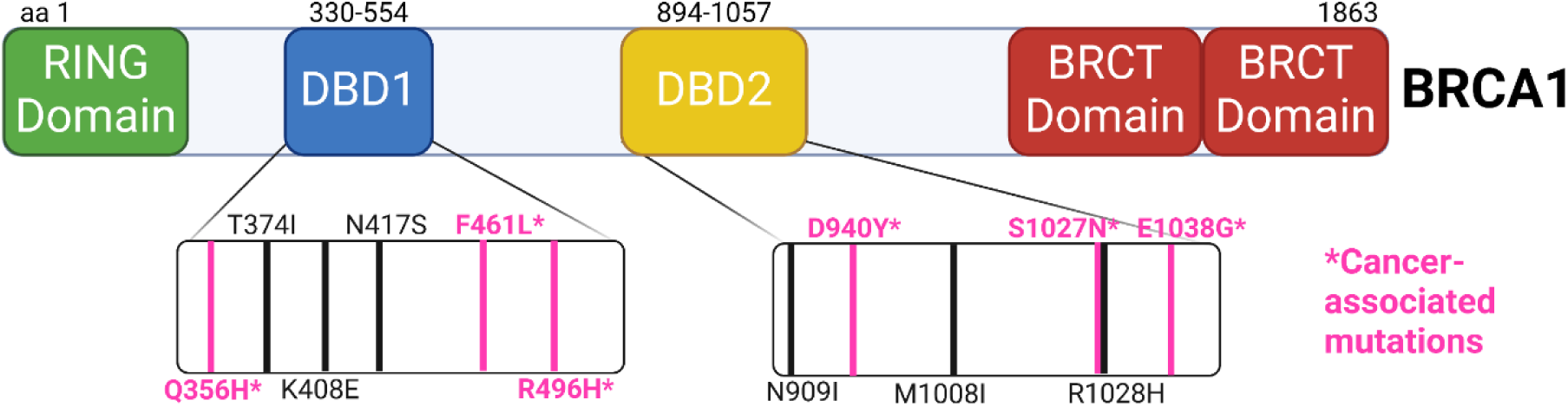
Map of BRCA1 depicting the approximate locations of the twelve DBD variants. Names of cancerous mutations are typed in pink. Names of VUSs are typed in black.

## MATERIALS AND METHODS

### Buffers and reagents

Buffers and reagents for site-directed mutagenesis, protein purification, and plasmid purification were prepared with research grade chemicals and Type I water that was purified from a Smart2Pure 6 UV/UF system (ThermoFisher; Waltham, MA, USA). Buffers and reagents for cell culturing and comet assays were prepared with research grade chemicals and Type I water that was purified from an Arium Mini Water Purification System (Sartorius AG; Göttingen, Germany). All solutions were sterilized through a 0.22 µm PES filter.

### Expression and purification of DBD constructs

*E. coli* expression constructs used to produce mutations for the BRCA1 DBD variants were purchased from Integrated DNA Technologies (Coralville, IA, USA). The primers in Table 1 were used to introduce mutations into the wild-type (WT) DBD constructs through site-directed PCR mutagenesis by use of a QuikChange II Site-Directed Mutagenesis kit from Agilent Technologies (Cedar Creek, TX). The resulting plasmids were sequence verified.

**Table 1.**
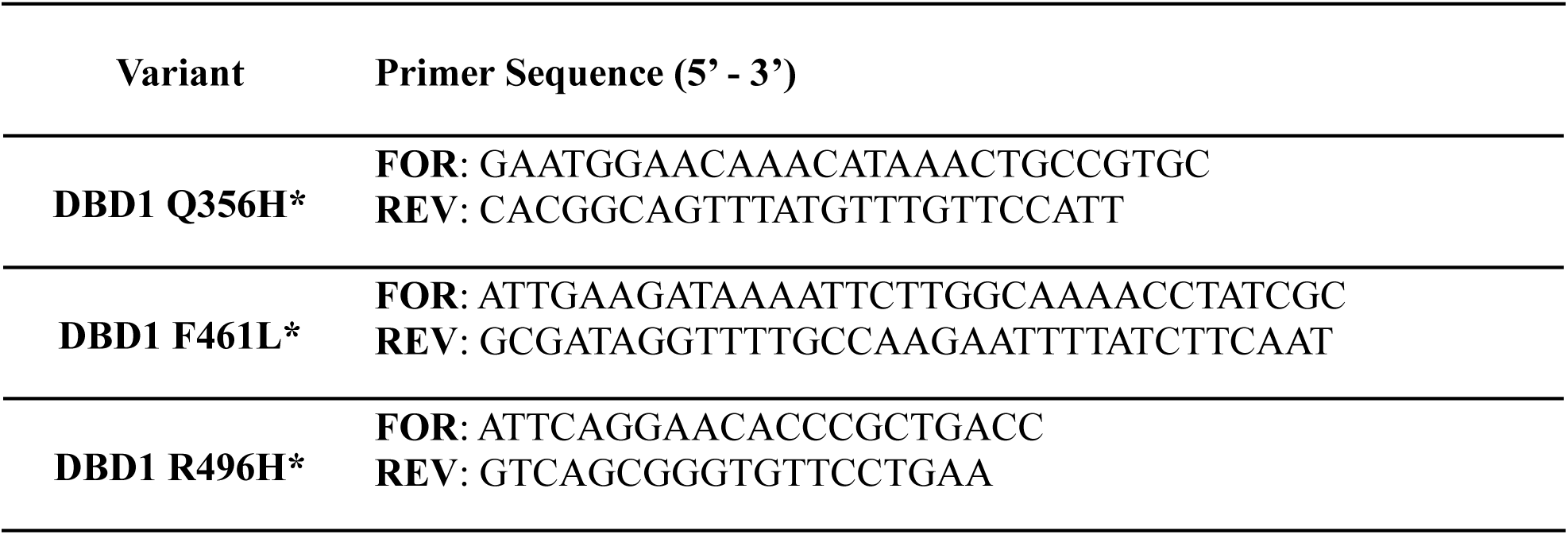
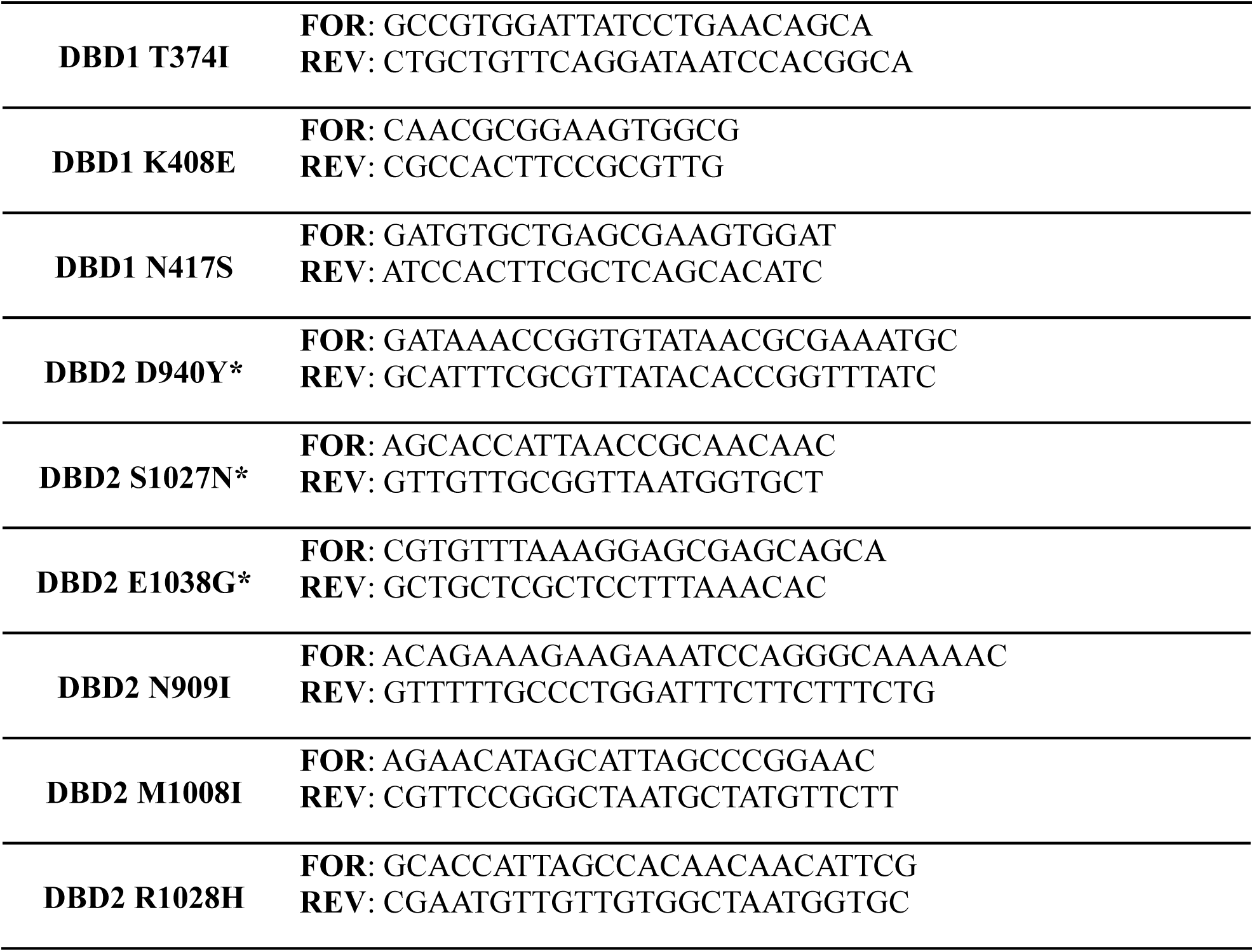
DNA primer sequences (*E. coli*). *Cancerous variants.

Plasmids were transformed into NiCo21(DE3) competent cells from New England Biolabs (Ipswich, MA, USA). 125 mL overnight LB growths were diluted into 4 × 1 L LB broth with 50 µg/mL carbenicillin and then incubated at 37 °C and 250 rpm using a MaxQ 4000 orbital shaker (ThermoFisher). DBD protein expression was induced with 1 mM IPTG upon reaching an OD_600_ of 0.6. Cells were harvested 4 hours post-induction by centrifugation using a J6-MI high-capacity centrifuge with a JS-4.2 rotor (4 °C, 4000 rpm, 90 min). The cell pellets were solubilized in lysis buffer (20 mM NaPi pH 7.5, 300 mM NaCl, 30 mM imidazole, 1 mM DTT, 5% glycerol, 1% NP-40). Sonication was performed on ice (15 seconds on, 45 seconds off, 45% amplitude for 30 minutes total) using a Fisherbrand 505 Dismembrator equipped with a 12 mm titanium probe (Fisher Scientific; Hampton, NH, USA). Lysates were clarified by centrifugation with a Sorvall RC5C Plus high-speed centrifuge and an SS-34 rotor (4 °C, 18,500 rpm, 90 min).

An Econo-Column (Bio-Rad; Hercules, CA, USA) was prepared with 20 mL of Ni-NTA agarose (Goldbio; St. Louis, MO, USA) and maintained at a temperature of 4 °C. Resin agarose was equilibrated with lysis buffer (20 mM NaPi pH 7.5, 300 mM NaCl, 30 mM imidazole, 1 mM DTT, 5% glycerol). Protein supernatant was filtered through a 0.45-micron PES filter and left to incubate with the resin for 30 minutes. The DBD proteins were eluted with elution buffer (20 mM NaPi pH 7.5, 30 mM NaCl, 250 mM imidazole, 1 mM DTT, 5% glycerol) and then loaded onto a 5 mL HiTrap Heparin HP column (GE Healthcare, Chicago, IL, USA) that was equilibrated with degassed binding buffer (1 mM DTT, 20 mM HEPES pH 7.5, 30 mM NaCl, 5 mM TCEP, 5% glycerol) using an AKTA Start FPLC system (GE Healthcare). The BRCA1 DBD proteins were eluted off the column with degassed elution buffer (1 mM DTT, 20 mM HEPES pH 7.5, 2 M NaCl, 5 mM TCEP, 5% glycerol) over a 50 mL linear gradient. Protein purity was assessed by SDS-PAGE. Fractions containing the BRCA1 DBD proteins were pooled and dialyzed with dialysis buffer (20 mM HEPES pH 7.5, 150 mM KCl, 1 mM DTT, 20% glycerol) in a 3500 MWCO Slide-A-Lyzer Dialysis Cassette. Protein concentrations were measured using a NanoDrop spectrophotometer (ThermoFisher) and known extinction coefficients at 280 nm, after which the proteins were snap-frozen in liquid nitrogen and stored at −80 °C for future use.

### DNA oligos

DNA substrates were created of single-stranded, double-stranded, and G-quadruplex DNA (ssDNA, dsDNA, and G4 DNA) for biolayer interferometry analyses. Oligos were synthesized by Integrated DNA Technologies (IDT) and are shown in Table 2. The complementary strand of the dsDNA was modified to include a Cy3 fluorescent label. Each substrate was annealed in G4 reaction buffer (20 mM HEPES pH 7.5, 150 mM KCl, 5 mM TCEP, 20% glycerol) through mixture of the top and bottom strands (in the case of dsDNA and G4DNA) with the non-biotinylated strand in molar excess of 1.05. Annealed samples were placed in a 95 °C heating block for 10 minutes and slowly cooled to 25 °C. Successful annealing was confirmed by native PAGE. G4 formation was separately validated by CD spectroscopy.

**Table 2.**
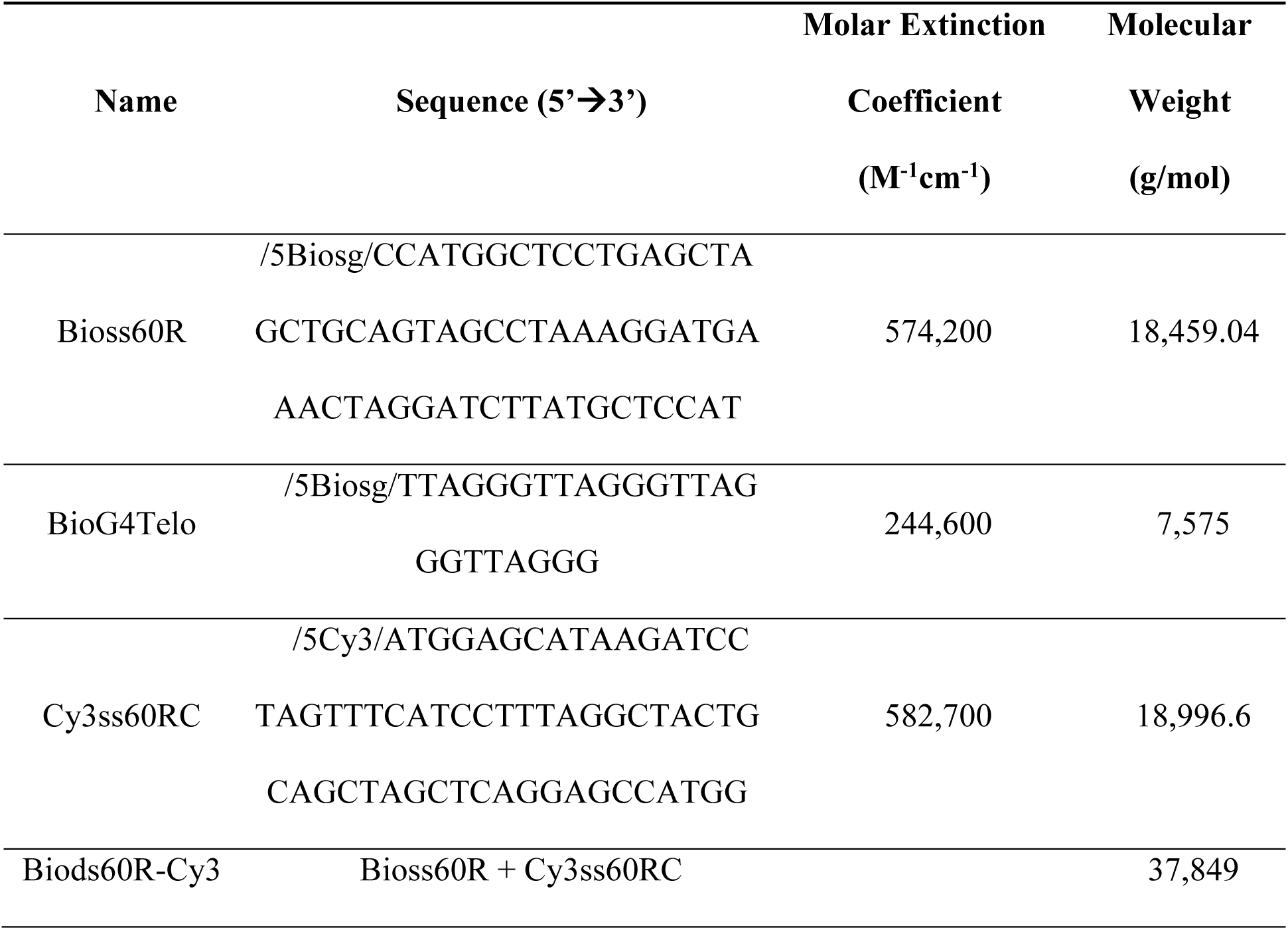
DNA sequences for biolayer interferometry analyses.

### Biolayer interferometry (BLI)

BLI studies were run on an Octet N1 instrument (Sartorius, formerly “BLItz”) and BLItz Pro instrument software in G4 reaction buffer (20 mM HEPES pH 7.5, 150 mM KCl, 5 mM TCEP, 20% glycerol) at 25 °C. High-precision streptavidin (SAX) biosensors (Sartorius) were hydrated 10 minutes in G4 reaction buffer prior to running the experiment. Samples were kept in Safe-Lock 0.5 mL amber microcentrifuge tubes (Eppendorf; Hamburg, Germany) at 25 °C and 250 rpm in an Eppendorf ThermoMixer before placement into the Octet N1 instrument for measurement. For each run, the interference pattern of the incident light was recorded in buffer alone for 30 seconds. The biotinylated DNA substrate was then introduced and allowed to bind to the biosensor through interactions with the streptavidin for 120 seconds. A wash was then performed in buffer alone for 30 seconds. Free BRCA1 DBD protein of varying concentrations was loaded and allowed to form protein-DNA complexes for 180 seconds. The biosensor was then returned to buffer for 120 seconds to observe dissociation of the protein-DNA complex. Runs were collected at each protein analyte concentration and concluded with a final reference in buffer alone to allow for comparison. Collection of BLI data was performed in triplicates for each DBD variant. For a detailed explanation of the methodology, please refer to our previous publication (41).

Binding isotherms were generated using the steady-state plateau values of the protein association phases plotted against total protein concentration. These values were fit to the simple one-site binding model in the following formula using GraphPad Prism software (Insight Partners; New York City, NY, USA), where the y-values represent the plateau signals, A is the amplitude, D is the DNA concentration (constant 150 nM), P is the BRCA1 DBD protein concentration, and K_D_ is the equilibrium dissociation constant (42). Average K_D_ and standard error values within a 95% confidence interval were reported for each variant.

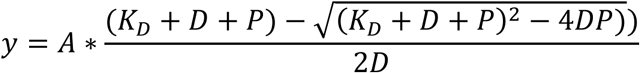

### Structural analyses

Predicted structures of the BRCA1 (UniProt: P38398) DBD proteins were virtually generated in Alphafold 2 (DeepMind; London, UK). Structural alignments were performed with PyMOL v2.5.4 (Schrödinger, Inc.; New York City, NY, USA).

### Plasmid purification of full-length BRCA1

The *E. coli* expression construct used to produce full-length human EGFP-BRCA1 was purchased from Addgene (Watertown, MA, USA) as a bacterial stab. Oligo primers were designed and obtained as described above. A 5 mL overnight starter culture was grown from the bacterial stab in LB broth with 5 µL carbenicillin, maintained at 37 °C and 250 rpm using a MaxQ 4000 orbital shaker (Thermo Fisher Scientific, Inc; Waltham, MA, USA). Plasmids were purified using the Nucleospin Plasmid miniprep kit and high-copy protocol by Macherey-Nagel Inc. (Allentown, PA, USA), then transformed into NEB 10-beta (DH10B) competent cells (New England Biolabs; Ipswich, MA, USA), from which 20% glycerol stocks were made. 5 mL overnight cultures were grown from the glycerol stocks and diluted into 250 mL of LB broth. Plasmids were then purified using the NucleoBond Xtra Midi Plus kit and high-copy protocol with NucleoBond Finalizer concentration (Macherey-Nagel). DNA concentrations were measured using a NanoDrop One C spectrophotometer (Thermo Fisher). Plasmids were sequence verified.

### Site-directed mutagenesis of full-length BRCA1

DNA oligonucleotide primers for site-directed mutagenesis were purchased from Integrated DNA Technologies (IDT; Coralville, IA, USA) at 100 µm concentrations. The primers in Table 4 were used to introduce mutations into the full-length BRCA1 construct. Site-directed mutagenesis was conducted using the QuikChange II Site-Directed Mutagenesis kit from Agilent Technologies, Inc. (Cedar Creek, TX). PCR products were digested with DpnI, then transformed into XL-1 Blue competent cells (Agilent) and NEB 5-alpha (DH5ɑ) competent cells (New England Biolabs). 5 mL cultures of the transformed colonies were grown and diluted into 250 mL of LB broth. Variant plasmids were then purified using the NucleoBond Xtra Midi Plus kit. DNA concentrations were measured using a NanoDrop spectrophotometer.

**Table 4.**
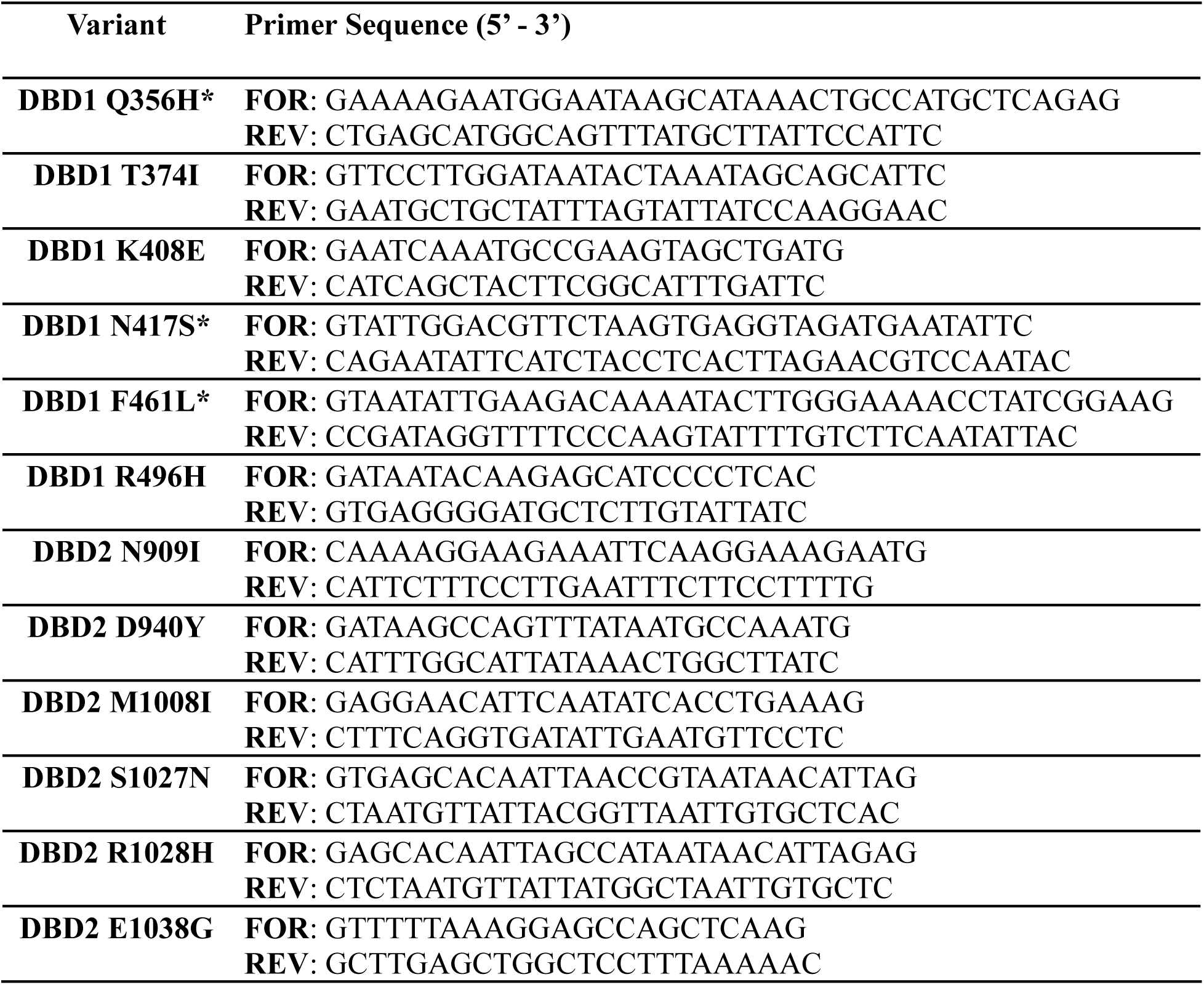
DNA primer sequences (full-length BRCA1). *Variants that did not undergo successful mutagenesis with the listed primers.

### Cell cultures and transfection

HEK293T cells were fed daily with DMEM/F12 1:1:Liquid (Hyclone Laboratories, Inc.; Logan, UT, USA) containing 10% FBS to passage 8, then frozen in a 5% DMSO solution and re-seeded two days before transfection. Transient transfection of wild-type and variant plasmids was conducted using Gibco Opti-MEM Reduced Serum Medium (Thermo Fisher) and 1% polyethylenimine (PEI) in DPBS (Hyclone). Cells were incubated in this solution for 24 hours before expression checks were conducted using a BioTek Cytation 1 Cell Imaging Multimode Reader (Agilent).

### Neutral comet assay and live cell counts

Cells were treated with bleomycin (Gold Biotechnology, Inc; Olivette, MO, USA) to induce DSBs at concentrations of 0, 1, 10, 50, 100, and 200 µg in DPBS for 15 minutes, or with menadione (Sigma-Aldrich; St. Louis, MO, USA) to induce oxidative stress lesions at concentrations of 0, 1, 10, 50, and 200 µM in DMEM for 1 hour. Cells were dissociated from growth vessels using 0.05% trypsin and neutralized with DMEM. Triplicate 20 µL samples of each treatment condition were taken and combined 1:1 with trypan blue for live-dead cell readings using a Countess II Automated Cell Counter (Thermo Fisher). The remaining cells were spun down at 700 rpm in an LC-8 centrifuge (Benchmark Scientific, Inc; Sayreville, NJ, USA) for 5 minutes, followed by aspiration of the supernatant and resuspension of the pellet in DPBS to a concentration of 5 × 10^5^ cells/mL. A solution of 1% Low Melt Agarose (Gold Biotechnology) in DPBS was combined in 500 µL portions with 50 µL aliquots of each cell solution, then spread on 2-well CometSlides (R&D Systems; Minneapolis, MN, USA) and refrigerated for 20 minutes to set. Slides were submerged in chilled CometAssay Lysis Solution (R&D Systems) for 1 hour. A solution of 10x neutral electrophoresis buffer was made with 1 M solid Tris and 3 M solid sodium acetate anhydride, then diluted 1:10. Slides were submerged in the 1x buffer for 30 minutes, followed by electrophoresis at 21 V in 1x buffer for 45 minutes using a Comet Assay Electrophoresis System II unit (Trevigen, Inc; Gaithersburg, MD, USA). Slides were immersed in a DNA precipitation solution of 1 M ammonium acetate in 95% EtOH at room temperature in the dark for 30 minutes, followed by 70% EtOH for 30 minutes, then dried overnight in a desiccator.

### Cell imaging and analysis

Comet slides were stained with 100 µL 0.001% SYBR Gold Nucleic Acid Gel Stain (Thermo Fisher) in DPBS per well for 30 minutes, then rinsed in Type I water and allowed to dry completely. Slides were imaged using Gen 5 BioTek Microplate Reader and Imager Software (Agilent) before analysis was performed using CometAssay Standardized System Comet Analysis Software (Trevigen). Between 100 and 200 individual cell images were used to take the mean value of the % DNA in tail and the comet moment for each treatment condition. A visualization of the general procedure is presented in Figure 3.

**Figure 3.**
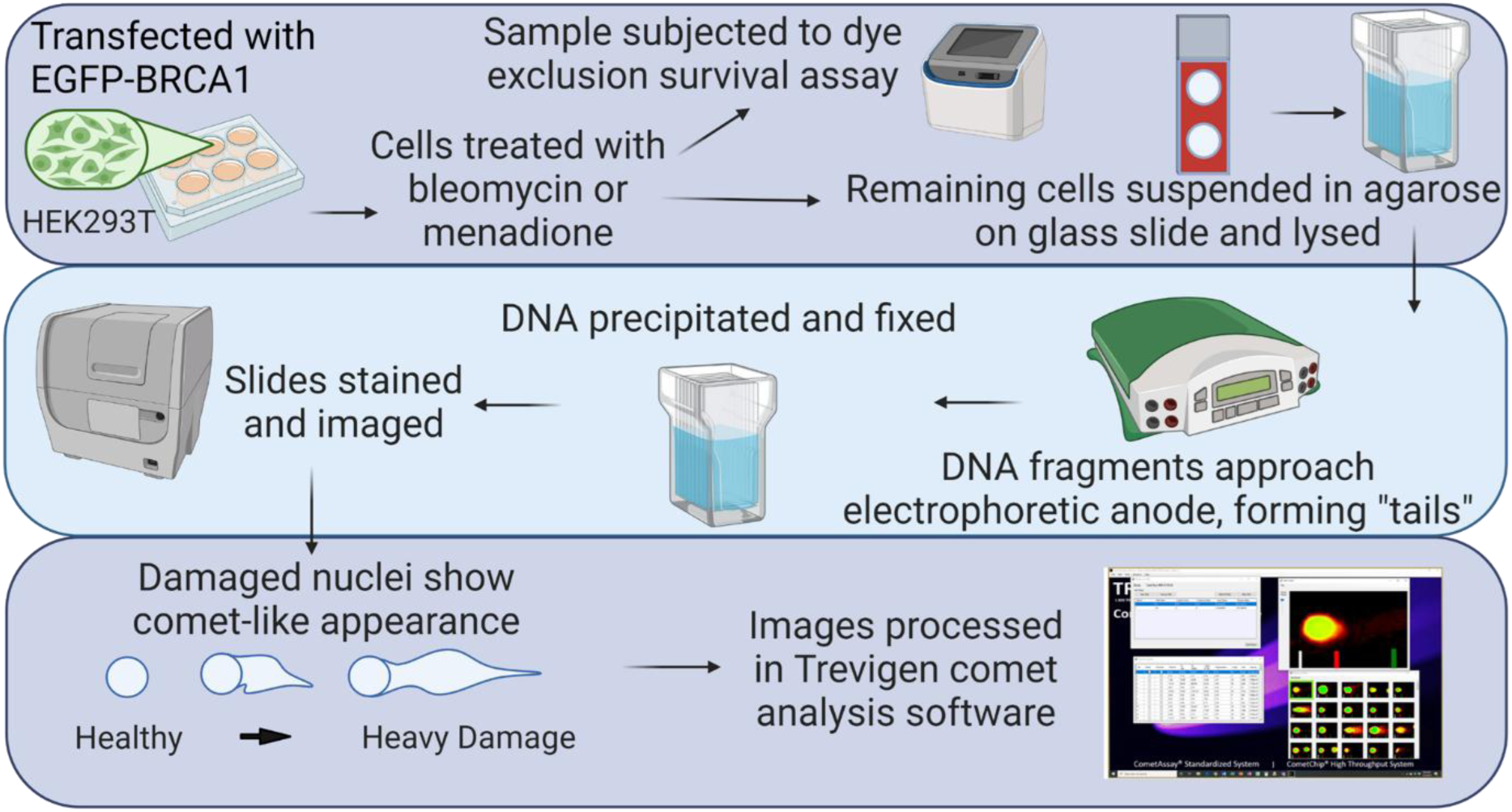
Visualization of neutral comet assay and cell imaging procedure.

## RESULTS

BRCA1 DBD variants were expressed and purified from *E. coli* as discussed in the Materials and Methods (Figure 4). Binding affinities were determined for each DBD variant with three different DNA repair intermediates (Table 2): single-stranded DNA (Bioss60R, or ssDNA), blunt-end double-stranded DNA (Biods60R-cy3, or dsDNA), and a G-quadruplex sequence (BioG4telo, or G4 DNA). The single-stranded and double-stranded sequences were studied as they appear as intermediates in the HR pathway. The G4 sequence appears in human telomeric DNA and was selected due to its involvement in BRCA1-mediated restart of stalled replication forks (43, 44). Protein-DNA binding interactions were determined using biolayer interferometry (BLI), a real-time label-free method for measuring biomolecular interactions (Figure 5).

**Figure 4.**
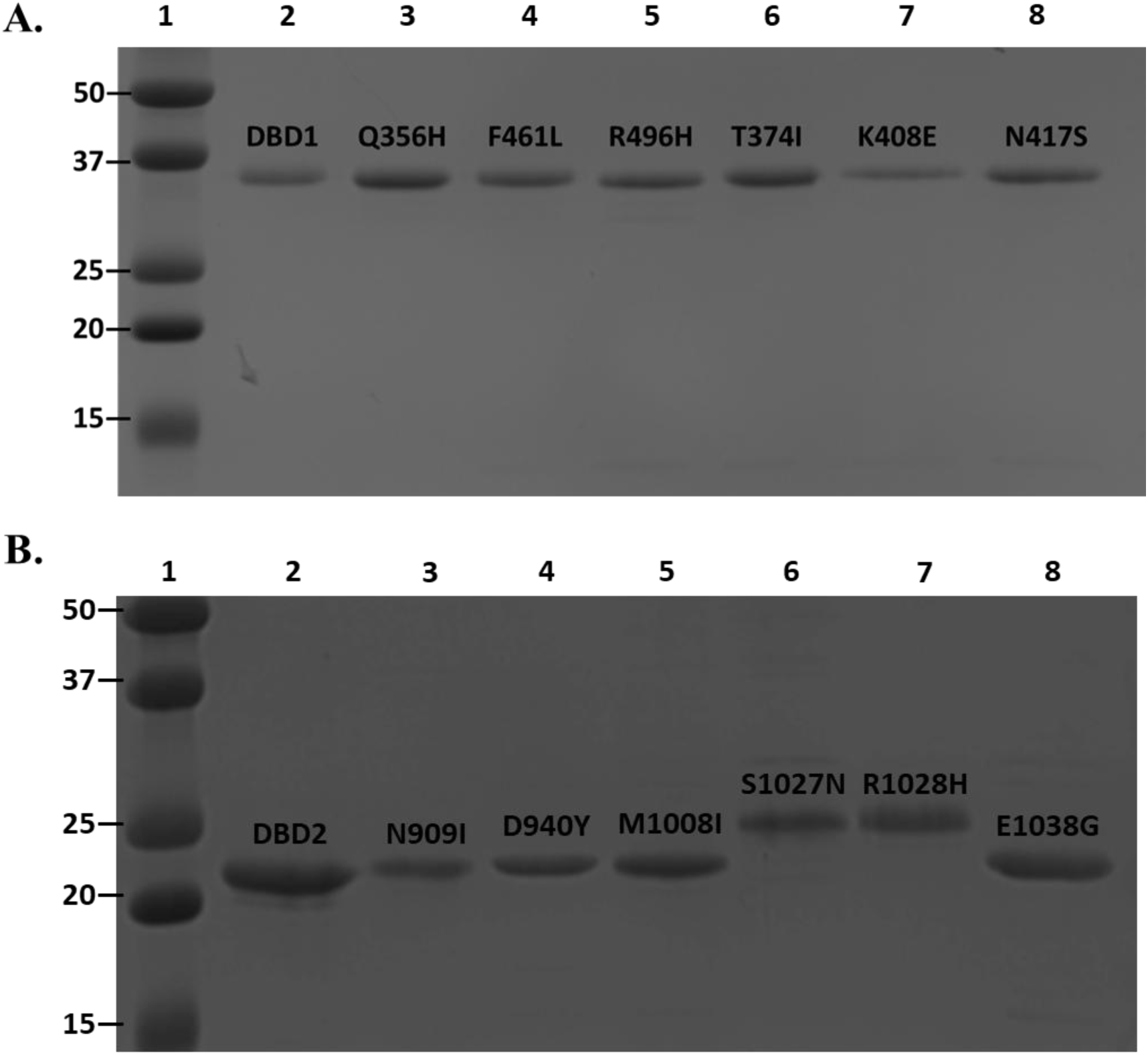
SDS-PAGE of purified recombinant proteins. **(A)** BRCA1 DBD1, DBD1 Q356H, DBD1 F461L, DBD1 R496H, DBD1 T374I, DBD1 K408E, and DBD1 N417S. Lane 1 is a protein ladder with known molecular weights (kDa), as labeled on the figure. Purified DBD1 was loaded in Lane 2 and resolved as a homogeneous band at ∼37 kDa. Purified DBD1 Q356H, F461L, R496H, T374I, K408E, and N417S were loaded in Lanes 3, 4, 5, 6, 7, and 8, respectively, and each migrated as ∼37 kDa bands. **(B)** BRCA1 DBD2, DBD2 N909I, D940Y, M1008I, S1027N, R1028H, and E1038G. Lane 1 is a protein ladder with labeled molecular weights in kDa. Purified DBD2 was loaded in Lane 2 and was observed at ∼22 kDa. Purified DBD2 N909I, D940Y, M1008I, S1027N, R1028H, and E1038G were loaded in Lanes 3, 4, 5, 6, 7, and 8, respectively, and each migrated as 22-25 kDa bands.

**Figure 5.**
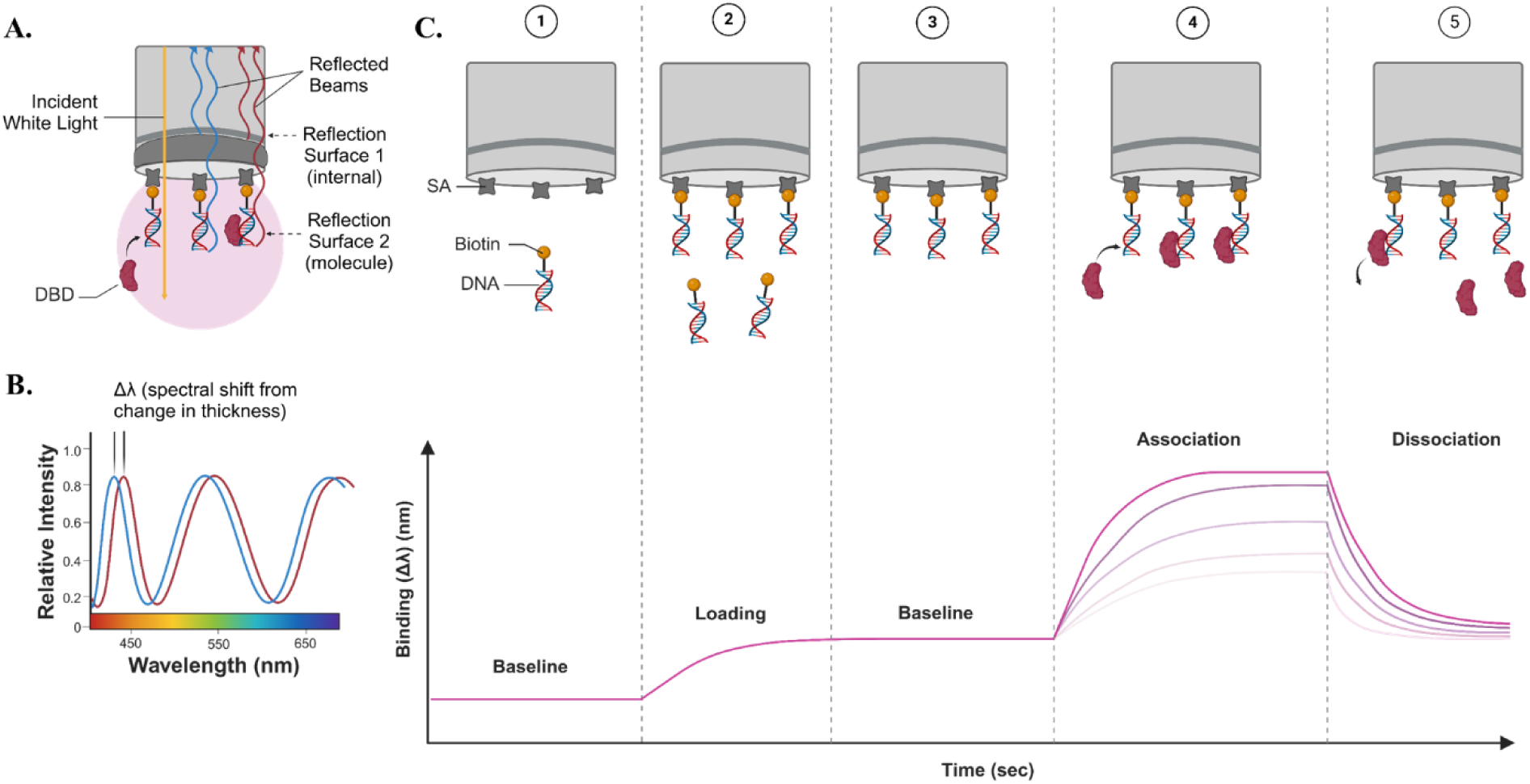
Biolayer interferometry (BLI) summary. (**A)** Graphic depicting the optical interference pattern of a high-precision streptavidin-coated (SAX) biosensor that allows the Octet N1 instrument to record protein-DNA interactions. The wavelength of the reflected beam differs based on what is bound to the surface of the biosensor. **(B)** Graphic depicting the spectral shift that occurs from a change in thickness. **(C)** Graphic depicting a time-course of the five steps in a typical BLI run. Phase 1 establishes an initial baseline in G4 reaction buffer alone. Phase 2 introduces biotin-tagged DNA that adheres to the SAX biosensor. Phase 3 reintroduces buffer to remove excess DNA that has not bound. Phase 4 introduces varying concentrations of the DBD proteins. As the proteins and DNA associate, the change in thickness is recorded as a spectral shift of the incident white light. Phase 5 reintroduces a final round of buffer which allows for the dissociation of the protein and DNA. The binding equilibrium plateau at the end of the association step (Phase 4) is plotted against protein concentration and fit to a 1:1 interaction model. A non-linear least squares analysis is used to calculate equilibrium K values.

The binding affinities of WT DBD1 for ssDNA, dsDNA, and G4 DNA were determined in a previous study and are included here to allow for direct comparison to the DBD1 variants (Table 4; Figure 6) (35). This work showed that WT DBD1 bound strongest to dsDNA (K = 2.92 ± 0.66 × 10^7^) and weakest to G4 DNA (K = 1.52 ± 0.28 × 10^7^). DBD1 Q356H had the same affinity pattern as WT DBD1, binding with the highest affinity for dsDNA (K = 8.49 ± 0.65 × 10^8^) and the lowest affinity for G4 DNA (K = 5.61 ± 2.15 × 10^7^). Notably, DBD1 Q356H bound to both dsDNA and ssDNA with a binding affinity that was an order of magnitude higher than seen in WT DBD1. DBD1 F461L also had the same affinity pattern as WT DBD1, with the strongest binding to dsDNA (K = 1.93 ± 0.55 × 10^7^) and the weakest binding to G4 DNA by an order of magnitude less (K = 1.33 ± 0.39 × 10^6^). DBD1 R496H bound strongest to ssDNA (K = 3.30 ± 0.79 × 10^7^) and weakest to G4 DNA by an order of magnitude less (K = 7.68 ± 1.20 × 10^6^). These results agree with cell phenotypes, which showed that cells overexpressed with the R496H variant demonstrated susceptibility to menadione compared to WT BRCA1 (Figure 7B, p < 0.05 in a T-test conducted using comet moment values from comet assaying under treatment conditions of 0, 1, and 100 μM menadione). Data for DBD1 Q356H and F461L remains to be gathered.

**Figure 6.**
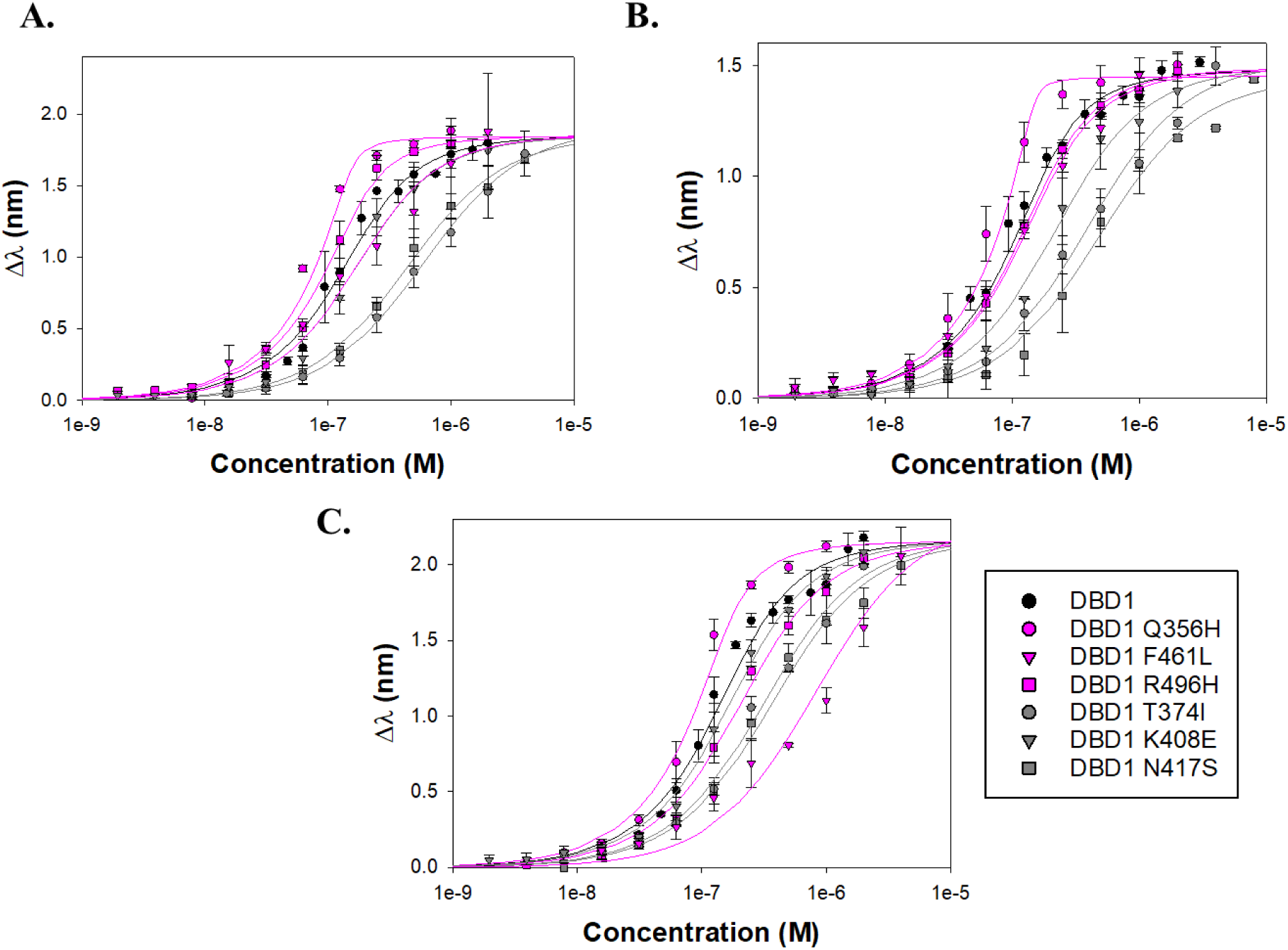
DNA binding affinity plots of BRCA1 DBD1 and variants. **(A)** ssDNA. **(B)** dsDNA. **(C)** G4 DNA.

**Figure 7.**
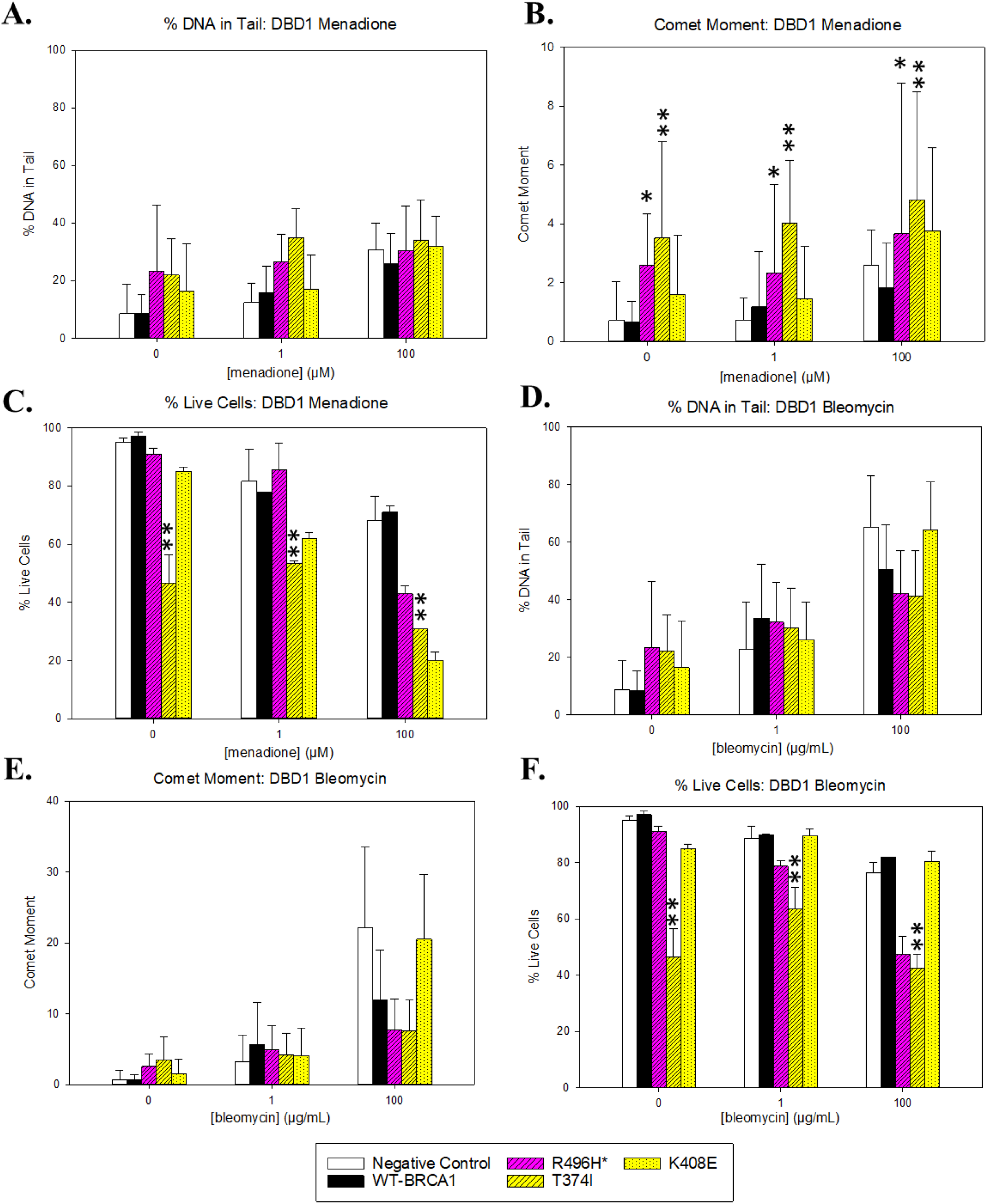
Cellular assays of BRCA1 DBD1. **(A)** Average percent of DNA in comet tails for DBD1 variants following tratment with menadione. None displayed significant differences from WT BRCA1. **(B)** Average comet moment for DBD1 variants following treatment with menadione. T374I and R496H showed significant increases in DNA damage compared to WT BRCA1 (p < 0.01 and p < 0.05 respectively). **(C)** Average percent live cells for DBD1 variants following treatment with menadione. Unclassified variant T374I showed a significant decrease in cell survival compared to WT BRCA1 (p < 0.01). **(D)** Average percent of DNA in comet tails for DBD1 variants following treatment with bleomycin. None displayed significant differences from WT BRCA1. **(E)** Average comet moment for DBD1 variants following treatment with bleomycin. None displayed significant differencese from WT BRCA1. **(F)** Average percent live cells for DBD1 variants following treatment with bleomycin. Unclassified variant T374I showed a significant decrease in cell survival compared to WT BRCA1 (p < 0.01). *Denotes known pathogenic variant.

The three unclassified VUSs showed an overall preference for G4 DNA (Table 4; Figure 6). DBD1 T374I had a distinct pattern of affinity and bound to all DNA repair intermediates with an order of magnitude lower than WT DBD1, with the strongest binding to G4 DNA (K = 4.12 ± 0.69 × 10^6^) and the weakest binding to ssDNA (K = 1.92 ± 0.13 × 10^6^). DBD1 K408E bound with similar affinity to ssDNA and G4 DNA (K = 1.06 ± 0.22 × 10^7^ and K = 1.07 ± 0.13 × 10^7^, respectively), and with the lowest affinity to dsDNA (K = 7.09 ± 1.09 × 10^6^). DBD1 N417S had a reversed pattern of affinity when compared to WT DBD1, binding strongest to G4 DNA (K = 3.42 ± 0.61 × 10^6^) and weakest to dsDNA (K = 2.48 ± 0.35 × 10^6^), with all binding affinities an order of magnitude lower than WT DBD1. Phenotypically, T374I had significantly decreased cell survival under both menadione and bleomycin treatment (Figures 7C and 7F). The variant also showed susceptibility to menadione by measurement of comet moment (Figure 7B). K408E, however, showed no significant difference from the wild-type protein under any condition (Figure 7). Cell survival data and DNA damage for cells overexpressed with the N417S variant remain to be quantified.

**Table 4.** Summary of BRCA1 DBD1 BLI data.

| Protein | DNA Sequence | K ($M^{-1}$) | A | R<sup>2</sup> |
| --- | --- | --- | --- | --- |
| **DBD1** | bioSS60R | $1.67 \pm 0.65 \times 10^7$ | $1.85 \pm 0.037$ | 0.995 |
| | bioG4telo | $1.52 \pm 0.28 \times 10^7$ | $2.16 \pm 0.044$ | 0.995 |
| | biods60R-cy3 | $2.92 \pm 0.66 \times 10^7$ | $1.48 \pm 0.021$ | 0.997 |
| **DBD1 Q356H\*** | bioSS60R | $2.97 \pm 0.61 \times 10^8$ | $1.65 \pm 0.053$ | 0.994 |
| | bioG4telo | $5.61 \pm 2.15 \times 10^7$ | $1.68 \pm 0.047$ | 0.995 |
| | biods60R-cy3 | $8.49 \pm 0.65 \times 10^8$ | $1.45 \pm 0.041$ | 0.997 |
| **DBD1 F461L\*** | bioSS60R | $1.06 \pm 0.41 \times 10^7$ | $1.98 \pm 0.144$ | 0.989 |
| | bioG4telo | $1.33 \pm 0.39 \times 10^6$ | $4.46 \pm 0.440$ | 0.986 |
| | biods60R-cy3 | $1.93 \pm 0.55 \times 10^7$ | $1.60 \pm 0.075$ | 0.995 |
| **DBD1 R496H\*** | bioSS60R | $3.30 \pm 0.79 \times 10^7$ | $1.51 \pm 0.024$ | 0.997 |
| | bioG4telo | $7.68 \pm 1.20 \times 10^6$ | $1.98 \pm 0.066$ | 0.997 |
| | biods60R-cy3 | $2.18 \pm 0.18 \times 10^7$ | $1.49 \pm 0.010$ | 0.999 |
| **DBD1**<br>**T374I** | bioSS60R | $1.92 \pm 0.13 \times 10^6$ | $2.10 \pm 0.133$ | 0.999 |
| | bioG4telo | $4.12 \pm 0.69 \times 10^6$ | $2.39 \pm 0.109$ | 0.996 |
| | biods60R-cy3 | $3.21 \pm 0.46 \times 10^6$ | $1.98 \pm 0.072$ | 0.996 |
| **DBD1**<br>**K408E** | bioSS60R | $1.06 \pm 0.22 \times 10^7$ | $1.75 \pm 0.067$ | 0.996 |
| | bioG4telo | $1.07 \pm 0.13 \times 10^7$ | $1.91 \pm 0.044$ | 0.998 |
| | biods60R-cy3 | $7.09 \pm 1.09 \times 10^6$ | $1.75 \pm 0.059$ | 0.997 |
| **DBD1**<br>**N417S** | bioSS60R | $2.74 \pm 0.24 \times 10^6$ | $1.72 \pm 0.023$ | 0.998 |
| | bioG4telo | $3.42 \pm 0.61 \times 10^6$ | $1.65 \pm 0.031$ | 0.997 |
| | biods60R-cy3 | $2.48 \pm 0.35 \times 10^6$ | $1.68 \pm 0.055$ | 0.996 |

Predicted structural models of WT DBD1 and the six variants were generated using AlphaFold 2 software and were used to compare relative positioning of residues. While WT DBD1 is predicted to be largely unstructured, it consists of three regions of alpha helices at residues 371-392, 457-461, and 514-517. The native Q356 residue is predicted to be located within a disordered region; the cancerous Q356H variant displays a rotated positioning of the histidine residue which now orients the residue towards a nearby alpha helix rather than outwards towards a solvent-accessible surface (Figure 8A). The cancerous F461L variant, located in the 457-461 alpha helix, has no altered orientation in comparison to the F461 residue (Figure 8B). R496H, also cancerous, experienced a minor change in the orientation of the histidine residue relative to R496 (Figure 8C). The T374I residue, located in the 371-392 alpha helix, is predicted to have no change in orientation from T374 (Figure 8D). Variants K408E and N417S (Figures 8E and 8F, respectively) experienced slight shifts in the positioning of the disordered loops, yet residues maintained similar orientation relative to the WT DBD1 predicted structure.

**Figure 8.**
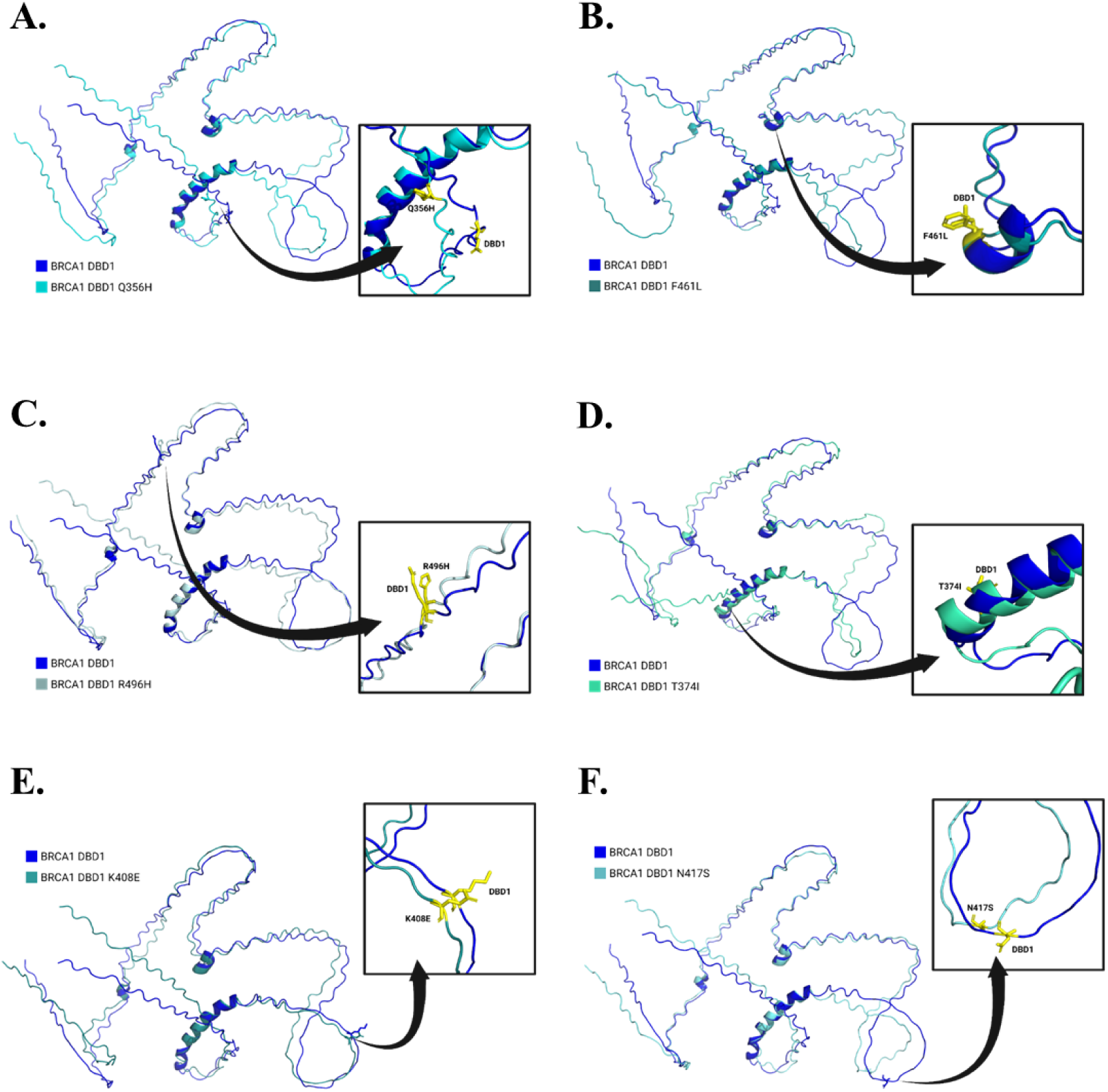
Structural analyses of the BRCA1 DBD1 proteins. **(A)** WT DBD1 and DBD1 Q356H. **(B)** WT DBD1 and DBD1 F461L. **(C)** WT DBD1 and DBD1 R496H. **(D)** WT DBD1 and DBD1 T374I. **(E)** WT DBD1 and DBD1 K408E. **(F)** WT DBD1 and DBD1 N417S.

As with WT DBD1, the binding affinities of WT DBD2 for ssDNA, dsDNA, and G4 DNA were determined in our previous work (35) and are included here for direct comparison to the DBD2 variants (Table 5; Figure 9). WT DBD2 bound strongest to G4 DNA (K = 3.26 ± 1.21 × 10^7^) and weakest to dsDNA (K = 2.27 ± 0.31 × 10^7^). All three DBD2 variants that are classified as pathogenic had DNA binding affinities that were lower than WT DBD2. DBD2 D940Y followed the same pattern of affinity as WT DBD2, preferring to bind G4 DNA most (K = 1.73 ± 0.08 × 10^5^) and dsDNA the least (K = 1.62 ± 0.28 × 10^5^); however, all DNA binding affinities were two orders of magnitude weaker. DBD2 S1027N, instead, bound strongest to ssDNA (K = 4.55 ± 0.23 × 10^4^) and weakest to G4 DNA (K = 1.09 ± 0.08 × 10^4^). DBD2 E1038G had the highest affinity for ssDNA (K = 9.80 ± 3.40 × 10^4^) and the weakest for dsDNA (K = 4.87 ± 0.49 × 10^4^). The DNA binding affinities of variants S1027N and E1038G were significantly weaker than WT DBD2, binding with strengths that are three orders of magnitude less. In cell-based assays (Figure 10), all three pathogenic DBD2 variants displayed differences from WT BRCA1 under at least one measurement condition. D940Y showed significant damage by measurement of comet moment under menadione treatment, indicating susceptibility to the oxidative stress-inducing agent. S1027N, too, showed susceptibility to menadione through a decrease in cell survival and increased comet moment compared to wild-type BRCA1. E1038G gave similar results, with significant DNA damage by measurement of both percent DNA in comet tail and comet moment.

**Figure 9.**
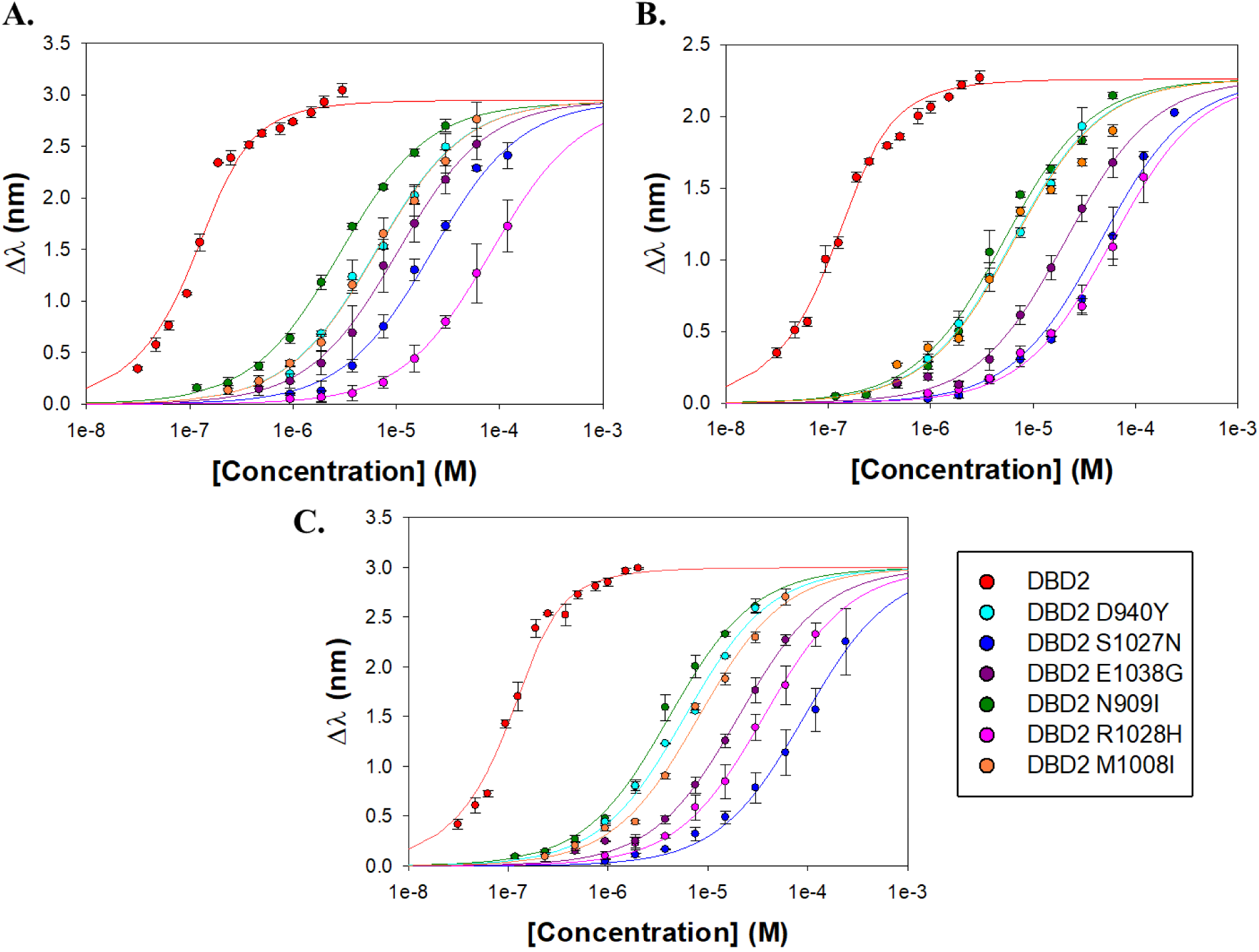
DNA binding affinity plots of BRCA1 DBD2 and variants. **(A)** ssDNA. **(B)** dsDNA. **(C)** G4 DNA.

**Figure 10.**
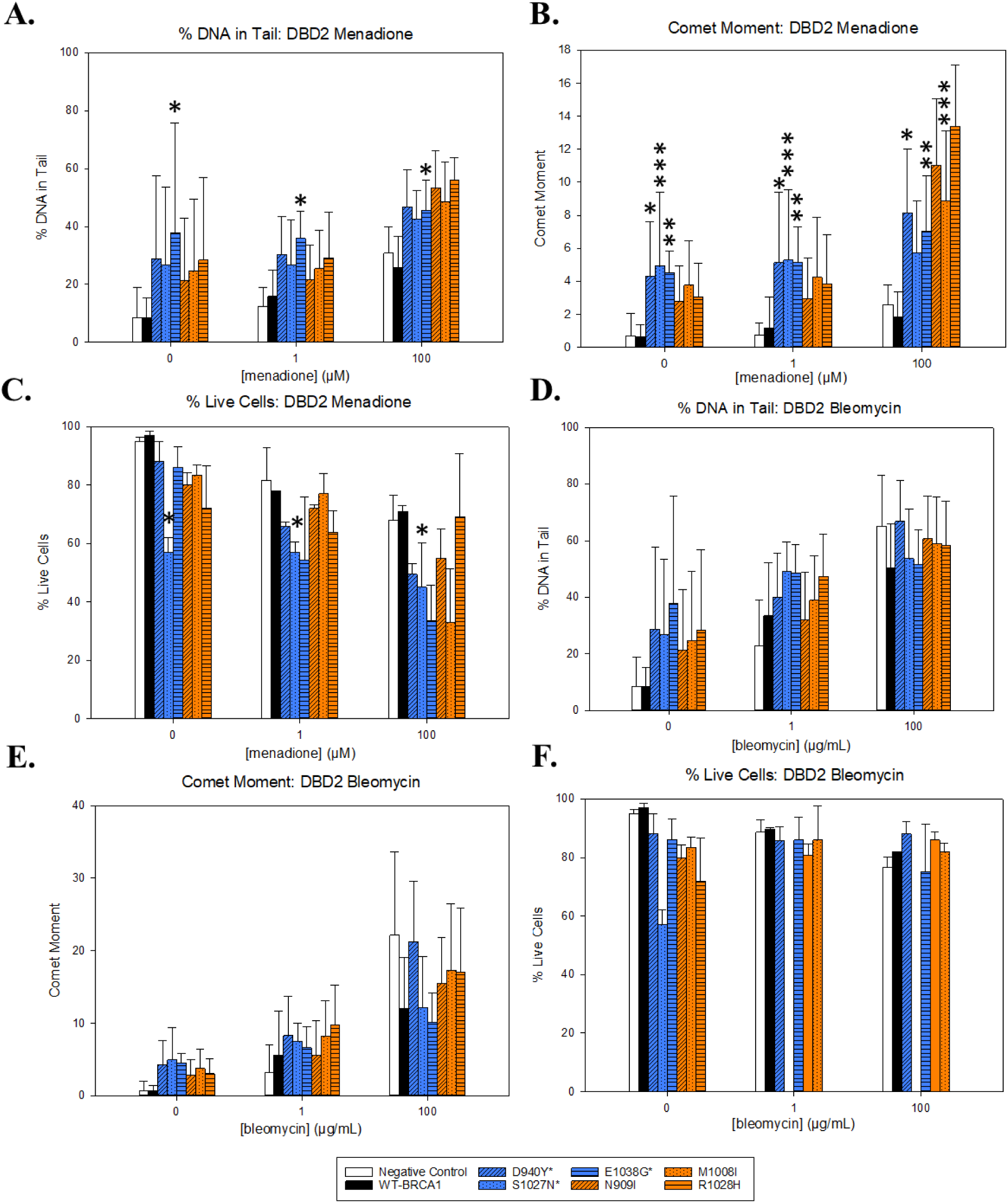
Cellular assays of BRCA1 DBD2. **(A)** Average percent of DNA in comet tails for DBD2 variants following treatment with menadione. Known pathogenic variant E1038G showed a significant increase in DNA damage compared to WT BRCA1 (p < 0.05). **(B)** Average comet moment for DBD2 variants following treatment with menadione. D940Y, S1027N, and E1038G showed significant increases in DNA damage compared to WT BRCA1 (p < 0.05, p < 0.001, and p < 0.01 respectively). **(C)** Average percent live cells for DBD2 variants following treatment with menadione. Known pathogenic variant S1027N showed a significant decrease in cell survival compared to WT BRCA1 (p < 0.05). **(D)** Average percent of DNA in comet tails for DBD2 variants following treatment with bleomycin. None displayed significant differences from WT BRCA1. **(B)** Average comet moment for DBD2 variants following treatment with bleomycin. None displayed significant differences from WT BRCA1. **(F)** Average percent live cells for DBD2 variants following treatment with bleomycin. None displayed significant differences from WT BRCA1. *Denotes known pathogenic variant.

**Table 5.**
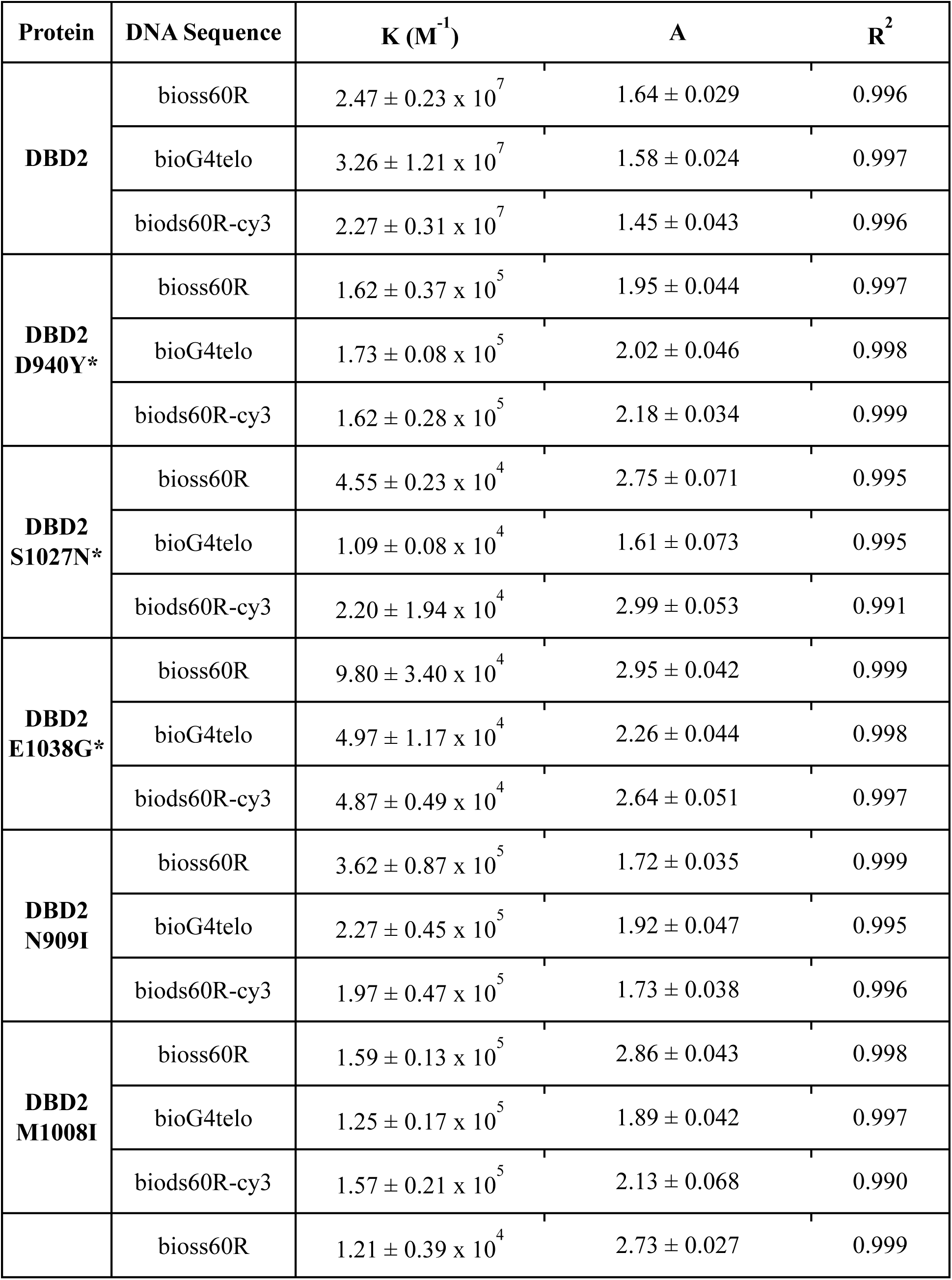
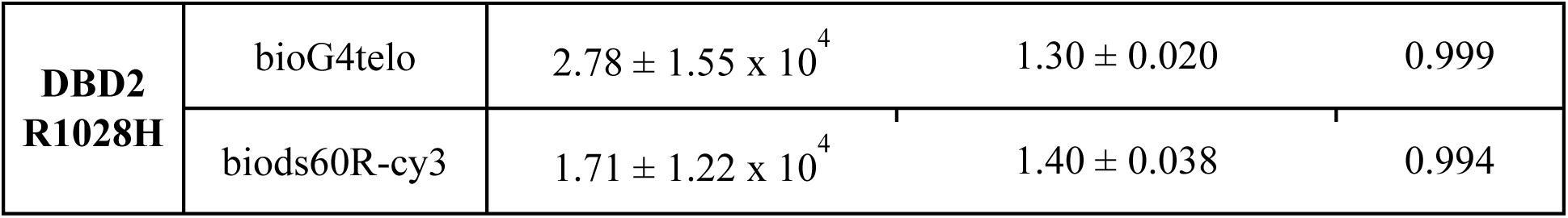
Summary of BRCA1 DBD2 BLI data.

The DNA binding affinities of the three unclassified DBD2 VUSs were either two or three orders of magnitude lower than those of WT DBD2. DBD2 N909I preferentially bound ssDNA (K = 3.62 ± 0.87 × 10^5^) and preferred dsDNA the least (K = 1.97 ± 0.47 × 10^5^). DBD2 M1008I bound strongest to ssDNA (K = 1.59 ± 0.13 × 10^5^) and weakest to G4 DNA (K = 1.25 ± 0.17 × 10^5^). DBD2 R1028H had the highest binding affinity for dsDNA (K = 1.71 ± 1.22 × 10^4^) and the weakest for ssDNA (K = 1.21 ± 0.39 × 10^4^). The unclassified DBD2 variants (N909I, M1008I, and R1028H) showed no significant differences from WT BRCA1 under any condition in the cell-based assays (Figure 10).

DBD2 variant AlphaFold models were generated and aligned to WT DBD2 in PyMOL. WT DBD2 is entirely unstructured with no secondary structures present; thus, we attempted to align structures to their first and last 40 residues. The cancerous D940Y variant displayed an altered position of the tyrosine residue in the variant, with the region surrounding the residue now facing inwards (Figure 11A). S1027N, a cancerous variant, had similar residue orientation but the region surrounding the residue was shifted (Figure 11B). The final cancerous variant, E1038G, displayed an altered residue positioning. Rather than facing downwards, the variant residue is facing in the opposite direction (Figure 11C). N909I and M1008I had slight shifts in the area surrounding residues, but both WT DBD2 and variant residues faced the same general direction (Figures 11D and 11E). R1028H showed a shifted loop and a small change in residue orientation (Figure 11F).

**Figure 11.**
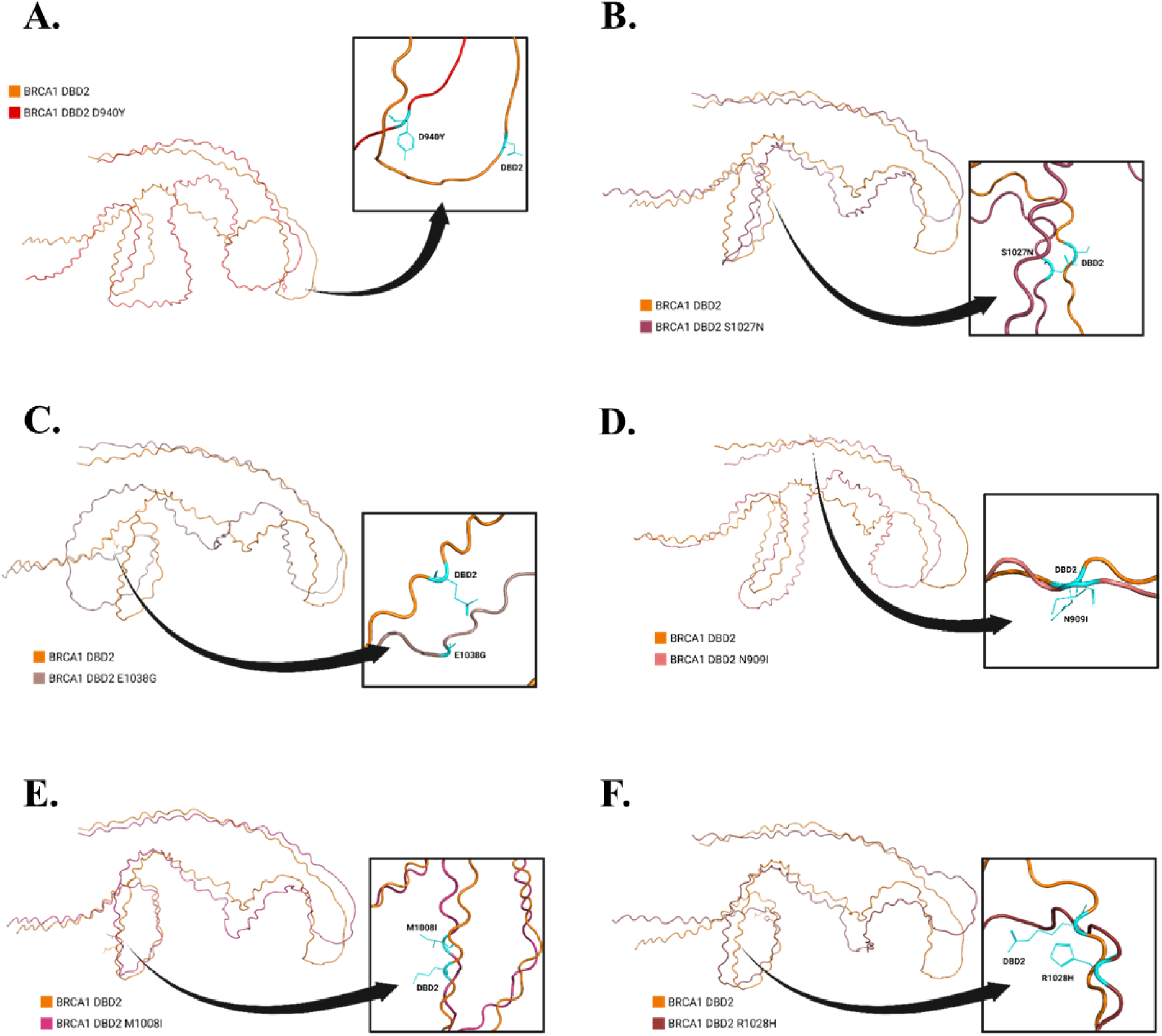
Structural analyses of the BRCA1 DBD2 proteins. **(A)** WT DBD2 and DBD2 D940Y. **(B)** WT DBD2 and DBD2 S1027N. **(C)** WT DBD2 and DBD2 E1038G. **(D)** WT DBD2 and DBD2 N909I. **(E)** WT DBD2 and DBD2 M1008I. **(F)** WT DBD2 and DBD2 R1028H.

## DISCUSSION

The central DBD of BRCA1, initially identified by Paull et al. as aa 452-1079, can be further studied as two separate domains: DBD1 and DBD2 (14). Based on other studies of DBD fragments (16, 17, 28, 30–32), we generated a BRCA1 DBD1 construct ranging from residues 330 to 554 and a BRCA1 DBD2 construct of residues 894 to 1057. The WT DBD constructs were studied in our previous work, which showed that WT DBD1 prefers to bind blunt-end dsDNA and may thus target BRCA1 to DSBs for their repair, while WT DBD2 binds strongest to ssDNA and G4 DNA and likely targets BRCA1 to telomeres for the repair of noncanonical structures (35). This study examined the DNA binding affinities of six pathogenic BRCA1 variants (Q356H, F461L, R496H, D940Y, S1027N, and E1038G) and six VUSs (T374I, K408E, N417S, N909I, M1008I, and R1028H) (36) for three DNA repair intermediates: dsDNA, ssDNA, and G4 DNA.

Computational research has previously classified half of the variants examined in this study as pathogenic, but these studies have not definitively classified the remaining variants. By comparing the unclassified BRCA1 variants to the binding specificities and cell phenotypes of WT BRCA1 and known cancerous variants, we can determine whether an unclassified point mutation may potentially lead to the development of cancer. While some variants maintained a similar pattern of affinity amongst the three DNA intermediates, most variants depicted an altered binding specificity that correlates to cellular susceptibility to menadione (oxidative stress) and structural changes that allow us to assign classifications to the unknown variants. The binding patterns, cell phenotypes, and computationally-predicted structures of the variants can also give insight to the functional roles of DBD1 and DBD2 in DNA repair pathways.

The BLI data of the three pathogenic DBD1 variants (Q356H, F461L, and R496H) reveals that each of these variants exhibited deviations from the DNA binding affinities and pattern of affinity of WT DBD1. For example, Q356H possessed binding affinities that were stronger for all DNA repair intermediates. Despite sharing the same pattern of affinity as WT DBD1, these affinities and the LOVD classification of Q356H as a cancerous variant (36) suggest that excessively strong binding to DNA may impede the ability of BRCA1 to accurately repair DNA. Interestingly, this may correspond with the gain in base stacking ability that occurs when Q356 is mutated to a histidine residue. Although this variant is predicted to reside within a disordered loop of DBD1, the mutation induces a significant shift in the spatial orientation of the residue and likely impacts its function. Cell data remains to be completed for this variant, but the determined binding characteristics and predicted structure support its pathogenicity and disruption of dsDNA repair.

As seen with Q356H, the cancerous F461L variant retained the same affinity pattern as WT DBD1. It possessed comparable affinities for dsDNA and ssDNA when compared to WT DBD1, yet had a reduced binding affinity for G4 DNA. Considering these results and the preference of WT DBD1 to bind dsDNA (35), one might initially conclude that this mutation has negligible effects on the DNA repair process as it did not experience a loss in dsDNA affinity. However, despite these findings, F461L is classified as a pathogenic variant by evolutionary and conservation analyses of mammalian *BRCA1*, which includes phylogenetic, ancestral sequence, and comparative evolutionary studies (45–47). The pathogenicity of F461L may be attributed to the substitution of an aromatic residue with an aliphatic one, which results in a loss of base stacking ability. While F461L retains its WT spatial orientation, the residue is situated within an alpha helix, and this loss in base stacking likely impacts the ability of DBD1 to effectively interact with G4 DNA. Cell data remains to be collected for this variant, but the alignment of its predicted structure to wild-type DBD1 shows displaced residues similarly to the other pathogenic DBD1 variants.

DBD1 R496H, also previously determined to be pathogenic due to co-occurrence with deleterious variants (48, 49), prefers ssDNA over other intermediates and exhibits susceptibility to menadione (oxidative stress). R496H displayed an overall preference for ssDNA, with a similar affinity for dsDNA and a much weaker affinity for G4 DNA. Structural predictions show that R496 is predicted to be located in a disordered loop and experiences minimal changes in spatial orientation when mutated to histidine. Unlike Q356H, which experienced an increase in DNA binding affinities when gaining base stacking abilities from a change to a histidine residue, R496H either maintained (as in the case of ssDNA and dsDNA) or saw a decrease (as with G4 DNA) in its binding affinities. Interestingly, comparative sequence, haplotype, trans-complementation, amino acid property-based conservation, evolutionary conservation, and biochemical scoring through A-GVGD analyses suggest that the variant is unlikely to be pathogenic (49–51). While these studies contrast our findings, both the DNA binding affinities and the phenotypic profile of this variant confirm its pathogenicity.

DBD1 T374I shows clear similarities to the pathogenic DBD1 variants. T374I demonstrated a preference for G4 DNA, bound an order of magnitude weaker to all DNA repair intermediates, and its overexpression in HEK cells results in significantly decreased cell survival and susceptibility to menadione. Predicted to be situated within an alpha helix, this mutation likely impairs DNA binding capabilities: through there exist minimal changes in the spatial orientation, we observe a slight bend of the helix following the mutated residue. Collectively, these results suggest that T374I may focus on targeting BRCA1 to non-canonical structures as opposed to DSBs. Thus, this VUS is likely pathogenic. This prediction is in alignment with evolutionary and conservation analyses of mammalian *BRCA1*, including phylogenetic, ancestral sequence, and comparative evolutionary studies, which classify T374I as a pathogenic variant (45–47).

K408E preferentially bound G4 DNA and ssDNA with similar affinities to WT DBD1, and with a diminished affinity for dsDNA. This suggests that K408E focuses primarily on targeting BRCA1 to non-canonical structures of DNA and may lead to pathogenicity; however, K408E exhibits no significant phenotypic differences in cells under the measured conditions, and the mutated residue is predicted to reside within a disordered loop which had minimal changes in spatial orientation upon mutation. As comparative sequence analyses of mammalian *BRCA1* suggest K408E to be a benign variant (50), the binding characteristics of K408E likely do not affect its DSB repair function; K408E is likely to be neutral.

N417S contrasted the affinity pattern of WT DBD1, preferring G4 DNA most and dsDNA least. Additionally, it bound with a weaker affinity – by an order of magnitude – to all DNA repair intermediates. N417S is predicted to reside within a disordered loop, yet the variant experienced a notable change in spatial orientation when mutated. Based on these results, N417S likely causes DBD1 to target BRCA1 to non-canonical structures of DNA. As these results contrast with comparative evolutionary studies and sequence-based predictions, which classify N417S as a benign variant (47, 50), it would be beneficial to conduct cellular studies of this variant as cellular data has not yet been collected.

The DBD2 variant D940Y is known to be pathogenic (47), bound G4 DNA two orders of magnitude less strongly than wild-type DBD2, and had susceptibility to menadione. Alignment of its predicted structure to the C and N termini of the predicted wild-type DBD2 structure results in significant displacement of residues throughout the domain and an altered orientation of residue 940. This structural difference likely results in the reduced DNA binding capability of the D940Y variant and impacts the ability of DBD2 D940Y to repair noncanonical DNA lesions arising from oxidative stress. The pathogenic variants DBD2 S1027N and E1038G (45, 46, 51) show similar patterns to each other, both binding ssDNA most strongly and displaying susceptibility to menadione. Like D940Y, their structural predictions differ throughout the entire domain from WT DBD2 when aligned, along with a changed spatial orientation of residue 1038 in the E1038G variant. These structural changes evidently result in decreased affinity for G4DNA, preventing the variants from addressing noncanonical structures as efficiently as WT DBD2.

DBD2 M1008I has a stronger affinity for ssDNA but exhibits no significant phenotypic differences from WT DBD2 in cells under the given conditions. Compared to the known pathogenic DBD2 variants, its predicted structure varies much less when aligned to WT DBD2. Evidently, the variant does not produce significant structural differences to alter binding affinity to such an extent that the noncanonical repair function of the domain is affected, even though the orientation of residue 1008 differs from WT DBD2. This is supported by computational studies that predict the M1008I variant to have a neutral effect (49–51). Previous studies involving DBD2 N909I are inconclusive (47); however, this variant exhibits the same patterns and characteristics as M1008I under the studied conditions, so it is likely that this variant is also neutral. R1028H, like WT DBD2, prefers G4 DNA and shows no significant phenotypic differences from the WT protein in cells. Like the DBD2 N909I and M1008I variants, its predicted structure shows less variation from WT DBD2 compared to the known pathogenic variants despite the altered orientation of residue 1028. Because this variant exhibits similar structural and binding characteristics to the wild-type domain, and has been previously predicted benign (50, 52), R1028H does not significantly disrupt the noncanonical repair function of DBD2.

None of the variants studied so far exhibit susceptibility to bleomycin (DSBs) except for DBD1 T374I by measurement of cell survival (Figure 7), suggesting that response to DSBs does not adequately predict pathogenicity of BRCA1 DBD variants. Other research argues that the mechanism resulting in cancer formation is ssDNA gaps that collapse into DSBs during DNA replication, which BRCA1 mutants then fail to repair (53, 54). The BRCA1 variants in this study could result in DNA damage by a mechanism not measured by the neutral comet assay. Additional studies should aim to study how ssDNA breaks affect BRCA1 variant responses in cells.

## CONCLUSIONS

Future studies should focus on study of variants Q356H, F461L, and N417S and their susceptibility to oxidative stress arising from bleomycin and menadione. As some of the variants showed susceptibility to menadione, likely as a result of the varied DNA lesions resulting from oxidative stress, it is possible that the treatment conditions used in this study were not adequate to incite a significant response of the BRCA1 variants to dsDNA breaks alone. Therefore, studies into other treatment conditions and DNA damage mechanisms may provide more insight into the variants that have inconclusive results. AlphaFold models are predicted models and cryo-EM should be used to verify structures.

## ASSOCIATED CONTENT

### ACCESSION CODES

BRCA1, UniProt: P38398

## Author contributions

C.G.W. designed experiments. E.C., E.L., V.S., K.L., and C.G.W. acquired data and interpreted results. E.C., E.L., V.S., K.L., and C.G.W. prepared the manuscript draft. All authors have given approval to the final version of the manuscript.

## ACKNOWLEDGEMENT

Figures 1-3, 5, 8, and 11, as well as the abstract graphic, were created with BioRender.com.

## ABBREVIATIONS

BRCA1: breast cancer susceptibility gene 1
HR: homologous recombination
DSB: double-strand break
DBD: DNA binding domain
aa: amino acid
BRCT: BRCA1 C-terminal repeats
ssDNA or Bioss60R: single-stranded DNA
dsDNA or Biods60R-Cy3: blunt-end double-stranded DNA
G4 DNA or BioG4Telo: G-quadruplex DNA
PALB2: partner and localizer of BRCA2
RAD51: DNA repair protein RAD51 homolog 1
RING: really interesting new gene finger domain
BLI: biolayer interferometry
SA: streptavidin

